# Inferring laminar origins of MEG signals with optically pumped magnetometers (OPMs): a simulation study

**DOI:** 10.1101/2023.08.20.554011

**Authors:** Saskia Helbling

## Abstract

We explore the potential of optically-pumped magnetometers (OPMs) to infer the laminar origins of neural activity non-invasively. OPM sensors can be positioned closer to the scalp than conventional cryogenic MEG sensors, opening an avenue to higher spatial resolution when combined with high-precision forward modelling. By simulating the forward model projection of single dipole sources onto OPM sensor arrays with varying sensor densities and measurement axes, and employing sparse source reconstruction approaches, we find that laminar inference with OPM arrays is possible at relatively low sensor counts at moderate to high signal-to-noise ratios (SNR). We observe improvements in laminar inference with increasing spatial sampling densities and number of measurement axes. Surprisingly, moving sensors closer to the scalp is less advantageous than anticipated - and even detrimental at high SNRs. Biases towards both the superficial and deep surfaces at very low SNRs and a notable bias towards the deep surface when combining empirical Bayesian beamformer (EBB) source reconstruction with a whole-brain analysis pose further challenges. Adequate SNR through appropriate trial numbers and shielding, as well as precise co-registration, is crucial for reliable laminar inference with OPMs.

## Introduction

Magnetoencephalography (MEG) is a non-invasive technique that measures the tiny magnetic fields generated by the synchronous current flow through neuronal populations in the brain^1,2^. Conventional MEG operates with superconducting SQUID magnetometers that must be immersed in liquid helium for cooling and are therefore placed within a fixed dewar for thermal insulation. This results in a substantial (several centimetre) gap between the sensors and the scalp and, as the strength of the magnetic field decreases with distance from the source, weaker magnetic fields at the sensor locations.

Optically-pumped magnetometers (OPMs) are highly sensitive magnetometers that operate without the need for cryogenic cooling. The sensors are small and lightweight and can therefore be flexibly arranged on, and placed close to, the scalp. OPMs contain helium gas^3^ or a vapour of alkali atoms^4–6^ whose atomic spins are aligned through optical pumping. Fluctuations in the local magnetic field affect the transmission of laser light through this spin-polarised gas or vapour and can be measured as changes in the amount of light at a photodetector site^7,8^. This new technology has sparked immense interest in the MEG community, and the first whole-head systems are now commercially available.

By placing the sensors directly on the scalp, the distance between the sensors and the cortical sources is decreased. Simulations show that bringing the sensors closer to the participant’s brain substantially improves the sensitivity to cortical sources and enables the sampling of higher spatial frequencies, i.e., this permits measurement of more focal field patterns, resulting in improved spatial resolution and better source separability compared to conventional MEG^7,9,10^. Experimental studies show that OPM-MEG systems exhibit comparable or even larger signal amplitudes and improved source reconstruction accuracy compared to conventional SQUID-MEG systems, even when they had fewer sensors with higher noise floors^11–14^.

A significant challenge in reconstructing the brain sources from MEG measurements taken on the scalp is to distinguish between sources originating from different cortical layers. Non-invasive laminar electrophysiology in humans would be of high relevance for both basic and clinical research, in particular in neurodegenerative diseases like Huntington’s disease^15^ and multiple sclerosis^16,17^ where different layers are affected across disease stages, and in investigating cortical microcircuits and inter-laminar neural communication^18^. However, with a cortical thickness of 2-5 mm^19,20^, the typical spatial resolution achieved by conventional MEG^21,22^ is not sufficient for laminar inference.

It is possible to distinguish between deep and superficial sources by using high-precision forward models^23–26^. Here, the main idea is to exploit the small variations in the so-called lead fields between deep and superficial sources to infer the more likely origin of the source activity, by comparing the model evidence of models that place the source activity in either the deep or the superficial layers. High-precision MEG has been used to accurately infer the origin of cortical dipole sources in simulations, applied experimentally to a visuo-motor paradigm^27^, and has been successfully used to derive laminar contributions to beta bursts in a temporally resolved manner^28^.

On-scalp OPM-MEG has been postulated to have the potential to further improve the discriminability of laminar sources^26,28,29^ as laminar inference was more successful for sources close to the sensors^26^. To investigate the conditions under which OPMs can improve the performance of laminar inference, we simulate cortical sources at deep and superficial layers and infer their laminar origin using high-precision forward models. We explore the impact of two main features of OPM sensor arrays: the number of sensors and the number of measurement axes. While a higher sensor density yields a higher spatial resolution in source reconstruction^7,9,30,31^, it raises questions of feasibility due to the potential for crosstalk and heating issues. OPM sensors with multiple measurement axes provide enhanced information capacity^10^ and have been shown to reduce external interference and motion artefacts substantially^32,33^. Additionally, the different sensitivity profiles of triaxial sensors allow for better and more uniform coverage, particularly for children^33^. We also investigate the effects of varying signal-to-noise ratios, sensor-scalp distances, co-registration errors, and interfering internal noise sources on the classification performance.

## Materials and Methods

Simulations were conducted using the development version of SPM (SPM12, release 6, https://github.com/spm/spm, downloaded 07/29/2022).

### MRI acquisition and processing

A structural MRI scan was used to reconstruct the laminar surfaces that define the source space of our simulations and to inform the source reconstruction forward model. We employed quantitative multi-parameter mapping data^34,35^ from a single participant (male, 23 years) from the MEG UK database (https://meguk.ac.uk/database). Acquisition was performed on a 3T Prisma scanner equipped with a 32-channel receive radio frequency (RF) head coil (Siemens Healthineers, Erlangen, Germany) and a body RF receive coil at the Wellcome Centre for Human Neuroimaging, UCL, London. The study was approved by the local ethics committee and the volunteer gave written informed consent before being scanned. The high resolution protocol was the same as in Bonaiuto et al. (2018)^26^, and consisted of three RF- and gradient-spoiled, multi-echo 3D FLASH scans with proton density-, relaxation time T1-, and magnetisation transfer-weighting (PDw, T1w, and MTw) at 800 μm isotropic resolution, plus a map of the RF transmit field B1 acquired using a 3D-EPI spin echo/stimulated echo method (SE/STE) corrected for geometric distortions due to spatial inhomogeneities in the static magnetic field B0^36^. For details on the acquisition protocol see Edwards et al. (2022)^37^.

Cortical surfaces were reconstructed using the recon-all pipeline from FreeSurfer (Fischl et al., 2004^38^; https://surfer.nmr.mgh.harvard.edu). Because the contrast in the quantitative MRI maps deviates significantly from the T1w MPRAGE image contrast expected by the recon-all pipeline^35^, the following steps were taken to extract an image with MPRAGE-like contrast from the 3T quantitative MRI parameters (see McColgan et al., 2021^39^). First, a small number of negative and very high values produced by estimation errors were set to zero in the longitudinal relaxation rate (R1) and PD maps, such that T1 (= 1/R1) was bounded between [0,8000] milliseconds (ms) and PD between [0,200] %. Then, the PD and T1 maps were used as input to the FreeSurfer mri_synthesize routine to create a synthetic FLASH volume with optimal white matter (WM)/grey matter (GM) contrast (repetition time 20 ms, flip angle 30°, echo time 2.5 ms). This synthetic image was given as input to the SPM segment function (https://www.fil.ion.ucl.ac.uk/spm) to create a combined GM/WM/cerebrospinal fluid (CSF) brain mask (threshold: tissue probability > 0), which was used for skull stripping. The skull-stripped synthetic image then served as input for the remaining steps of the recon-all pipeline which resulted in cortical surfaces for the GM/WM boundary (‘white’ surface), the pial surface and mid-cortical surface.

### Simulated OPM-MEG sensor arrays and data

We simulated single dipolar sources as measured by OPM-MEG sensor arrays of different configurations for all simulations. We simulated OPM datasets with 200 trials with a duration of 1000 ms and a sampling rate of 200 Hz using code adapted from https://github.com/tierneytim/OPM. Sensor locations were determined using a point packing algorithm that positions sensors on the scalp surface at increasing densities as described previously^40^. Each sensor was modelled with one to three measurement axes: single radial axis, radial axis and one transversal axis, radial axis and two orthogonal transverse axes. We considered array designs with sampling distances between 25 and 55 mm in 10 mm increments, corresponding to 32, 42, 71, 138 sensors for the single axis array configurations.

The sensor-scalp offset was set to 6.5 mm, following Tierney et al., 2020^40^. This corresponds to a typical distance between scalp and the centre of sensitivity of commercially available SERF OPM sensors (2nd generation QuSpin OPM sensors). We also simulated off-scalp MEG with increased scalp-sensor offsets of 20, 30 and 40 mm, using otherwise identical simulation parameters. Our aim was to compare laminar inference performance of OPM-MEG sensor arrays to conventional SQUID-based MEG, where sensors are placed further away from the scalp and scalp-sensor offsets in the range of 15 to 40 mm are typical^14,41–43^. Varying the scalp-sensor offsets also allowed us to evaluate the impact of possible design compromises concerning sensor placement in the OPM-MEG array, e.g., due to the use of generic helmets^44^ or the necessity to accommodate cooling pads in high-density sensor packaging in the case of alkali-based OPMs.

We simulated deep and superficial current dipole sources at vertices on the white matter/grey matter boundary surface and the pial surface meshes, respectively. The meshes were constructed using the cortical surfaces (’white’ and ‘pial’) derived from the FreeSurfer surface reconstruction pipeline applied to an individual’s structural MRI, as detailed in the “MRI Acquisition and Processing” section. The mesh resolution was determined by the pre-specified cortical meshes derived from the FreeSurfer surfaces, downsampled by a factor of 10. This resulted in 32,212 vertices for each surface, with an average vertex spacing of 1.74 mm for the white surface and 1.92 mm for the pial surface. Sources were positioned at mesh vertices, and the orientation of each cortical current source was defined by the surface normal of the cortical mesh at that location. The same meshes were later used for source-reconstruction (See “Laminar source estimation” section).

For each surface, we randomly selected 60 vertices as cortical source locations. At each source location, a 20 Hz sinusoidal dipolar source patch with a patch size of 5 mm at Full Width at Half Maximum (FWHM) was added for each of the 200 trials modelled per cortical source location. The dipolar sources were active for 400 ms. Synthetic datasets were generated across a range of realistic SNRs by adding Gaussian white noise to the simulated data, scaled to yield per-trial amplitude SNR levels (averaged over all sensors), of −50, −40, −30, −20, −10, −5 dB. The forward model linking the simulated dipole current source to the sensor level activity was based on the Nolte single shell approach with the inner skull surface and cortical surfaces derived from the structural MRI. Sensors were assumed to be point magnetometers.

### Laminar source estimation

We next aimed to determine the laminar origin of the simulated sources from the sensor data alone. We used two main types of analyses: a whole-brain and an ROI-based analysis, equivalent to those described in Bonaiuto et al., 2018^26^.

In the whole-brain analysis, we reconstruct the OPM-MEG sensor data once to the pial and once to the white matter cortical surface and then compare the fit of the two models using Bayesian model comparison as a metric^23,26^. For Bayesian model comparison, we compute the difference in free energy between the pial and white matter forward models, approximating the log ratio of the model likelihoods. This results in a metric that is positive or negative, if there is more evidence for the pial or white matter model, respectively. A difference in log model evidence greater than 3 indicates that one model is approximately 20 times more likely than the other.

While this whole-brain analysis provides a global answer to the question of which cortical surface the sensor activity is more likely to originate from, it lacks spatial specificity regarding the location within the cortex where this laminar activity originates. To address this limitation, a region-of-interest (ROI) analysis that reconstructs the data onto both pial and white matter surfaces simultaneously can be employed^26^. An ROI is calculated based on the change of activity on either surface from a baseline time window, and the reconstructed activity within the ROI is compared between the two surfaces. We functionally defined ROIs by comparing power in the 10–30 Hz frequency band during the time period containing the simulated activity ([100 500] ms) with a prior baseline period ([-500 100] ms) at each vertex using two-tailed paired t-tests. Vertices in either surface with a t-statistic in the 75th percentile of the t-statistics over all vertices in that surface, as well as the corresponding vertices in the other surface, were included in the ROI. For each trial, we computed ROI values for the pial and white matter surfaces by averaging the absolute value of the change in power compared to baseline in that surface within the ROI. Finally, we used a paired t-test with variance regularisation^45^ to compare the ROI values from the pial surface with those from the white matter surface over trials. The ROI analysis produced a t-statistic which was positive when the change in power was greater on the pial surface and negative when the change was greater on the white matter surface.

In both analyses, we estimated sources using the empirical Bayes beamformer (EBB^46,47^) and multiple sparse priors (MSP^48^) source reconstruction approaches as implemented in SPM12 (https://www.fil.ion.ucl.ac.uk/spm/). The corresponding functional priors assume a sparse distribution of current flow across the cortex, uncorrelated in time for the EBB and locally coherent and sparse for the MSP approach. As in Bonaiuto et al. (2018), for the whole brain analysis the source inversion was applied to a Hann windowed time window from 500 ms to −500 ms filtered from 10–30 Hz, while no Hann window was used for the ROI analysis. These data were projected into 274 orthogonal spatial (lead field) modes and 4 temporal modes. Cortical patches were modelled with a patch size of 5 mm FWHM. In this study, we used the development version of SPM as it calculates geodesic distances in the cortical mesh construction in an exact manner, while previous SPM implementations used the approximate Dijkstra algorithm.

We did not investigate simulations with minimum norm^2^ and LORETA^26^ source localisation, as they have been shown in the past to be unable to allow laminar inferences for simulated sparse sources^26^. A replication of these findings can be seen in Fig. S2 in the Supplementary material. We note that we simulated sparse sources and that our inability to distinguish between different laminar sources using the IID or LORETA approaches can be at least partly attributed to the mismatch between data and functional prior assumptions^23,26^.

The code for the current dipole sources simulations and the laminar inference was based on and adapted from the code at https://github.com/jbonaiuto/laminar_sim.

### Impact of co-registration errors

Laminar MEG using conventional SQUID-MEG relies on subject-specific head-casts to reach the required accuracy in the high-precision forward models. To build the forward models, the accurate position and orientation of MEG sensors relative to the cortical surfaces derived from the MRI (i.e., co-registration of the two modalities) needs to be established. Head-casts enable highly accurate co-registration in the sub-millimetre range and reduce head movements during scanning to less than 1 mm. This results in better data quality and anatomically more precise MEG recordings^24,25^ than for conventional co-registration strategies based on fiducials and surface mapping which typically achieve an accuracy of 5–10 mm^47,48^ (but see Sonntag et al. 2018^49^ who used an adaptive Metropolis algorithm to reach target registration errors between 1.3 and 2.3 mm at the head surface).

Accurate co-registration is also essential to fully utilise the high spatial resolution of on-scalp MEG systems. The expected co-registration error for OPM systems differs between flexible and more rigid sensor arrays^13^, with rigid helmets being preferred for high-precision measurements due to their more accurate estimation of relative sensor locations and orientations^14^. Here, we investigate the effect of co-registration errors expected for rigid sensor arrays by introducing random displacement to the three fiducial locations, with standard deviations from 1 to 4 mm in 1 mm increments. Note that the impact of potential movements of the participant’s head in relation to the rigid sensor array during the experiment was not investigated in our study and needs to be further investigated.

### Impact of source patch sizes

In the simulations described so far, we assumed cortical source patches with a width of 5 mm and used smoothed MSP and beamforming priors based on an equivalent smoothing kernel (as implemented in spm_eeg_invert_classic.m), i.e., the estimated patch extent in our source reconstructions fitted the true simulated source patch extent. Previous simulation studies^23,26^ have demonstrated that the under- or overestimation of patch extent can introduce biases in model evidence. Additionally, in experimental work^27^, it was observed that smaller source prior patch sizes introduced a superficial bias, while larger patch sizes introduced a deep laminar bias. We thus performed a set of simulations with congruent and incongruent patch sizes, using patch sizes of 5 and 10 mm, to investigate the impact of incongruent patch sizes on laminar inference performance for on-scalp OPM sensor arrays.

### Adding internal interference sources

While simulations of single current dipoles can be illustrative, they do not reflect real experimental situations where laminar inference is performed on a source of interest in the presence of other interfering brain sources. To investigate the impact of such internal interfering sources, we simulated additional noise sources on the mid cortical surface. The source time courses of the source of interest and of five internal noise sources were modelled as Gaussian random data within a frequency range of 10 to 30 Hz and the source amplitudes of the internal noise sources were defined in proportion to the source of interest with a relative source strength of 0.4.

### Statistics

We first compared the classification accuracy and bias of each analysis and source inversion algorithm by computing the percentage of sources that were classified correctly and the percentage of sources classified as coming from the pial surface and subjected these percentages to two-sided binomial tests with chance levels of 50% to evaluate their significance. To investigate whether these effects were also significant at the single simulation level, we additionally employed a threshold of ±3 for the free energy difference (meaning that one model is approximately twenty times more likely than the other) and a threshold of the critical t-value with degrees of freedom (df) = 199 and *α* = 0.05 for the ROI t-statistic. By doing so we examined whether the classification approach not only found a difference in the correct direction, but also whether this metric was significant.

We used logistic regression to evaluate changes in classification accuracy and bias across sampling densities, number of axes, co-registration errors, and sensor-scalp offsets. Differences in laminar inference performance for free energy and the ROI-t-statistic analysis and for congruent and incongruent patch sizes were evaluated using exact McNemar’s tests^49^.

## Results

### At which signal-to-noise ratios can we draw laminar inferences?

We first evaluate the classification accuracy and bias across SNRs for an OPM-MEG array with a 35 mm inter-sensor distance to examine at which SNRs laminar inferences can be made using non-invasive OPM-MEG. We also report differences in laminar inference performance between the whole-brain free energy and the ROI-t-statistic analysis. Results for the OPM sensor array with a 35 mm inter-sensor distance are summarised in Figure 1.

Classification accuracy improved with increasing SNR for both source reconstruction approaches and laminar inference analyses (EBB/free energy: beta = −1.168, p <.001; EBB/ROI-t-statistic: beta = −2.038; p <.001; MSP/free energy: beta = −2.941 p <.001; MSP/ROI-t-statistic: beta = −3.341, p <.001), while the percentage of sources classified as coming from the pial surface decreased with increasing SNR (EBB/free energy: beta = 1.292, p = <.0.001; EBB/ROI-t-statistic: beta = 1.855; p <.001; MSP/free energy: beta = 0.456, p <.001; MSP/ROI-t-statistic: beta = 0.350, p <.01. We note that for the EBB source reconstruction approach at high SNRs, classification was biased towards the deep surface when using the whole-brain analysis.

For the EBB source reconstruction approach, laminar source inference was statistically significant at SNRs of −20 dB or higher; we observed however a bias towards the deep surface for the free energy metric. At these relatively high SNRs, the ROI-based analysis performed better than the whole-brain analysis: performance accuracy was higher (two-sided exact McNemar’s tests: p <.001 at SNRs of −5, −10 and −20 dB) and classification less biased (two-sided exact McNemar’s tests: p <.01 at SNRs of −5 and −10 dB). At an SNR of −30 dB, the ROI approach failed to yield significant results at the single simulation level, i.e., the absolute t-statistic values did not exceed the significance threshold. Additionally, laminar inference was biased towards the pial surface. While the free energy metric yielded a larger ratio of simulations with statistically significant classifications, classification accuracy was low. No laminar inference was possible at a very low SNRs of −40 and −50 dB for both analyses, and classification accuracy was at chance level and classification strongly biased towards the pial surface.

The MSP approach yielded high classification performances and no significant biases at SNRs of −30 dB or higher. Note that, there was a high accordance between the simulated data and the prior assumptions of the MSP approach, rendering this approach somewhat idealised. Classification performance did not differ significantly between the ROI-based and the whole-brain analysis as shown by two-sided exact McNemar’s tests for classification accuracy and bias. At an SNR of −30 dB, however, the ROI approach failed to yield significant results reliably at the single simulation level. At very low SNRs of −40 and −50 dB, classification accuracies and biases were not statistically significant at the single simulation level for both, whole-brain and ROI-based, analyses.

In summary, we find that for an OPM-MEG sensor array with an inter-sensor distance of 35 mm using the MSP approach (with patch priors that included the source locations) allowed us to achieve highly accurate laminar inferences at SNRs of −30 dB or higher. For the EBB approach an SNR of at least −20 dB was required to reliably infer the correct laminar origin of simulated sources. Additionally, the ROI-based analysis is recommended over the whole-brain free energy analysis due to its superior classification accuracy and reduced bias.

**Fig. 1.**
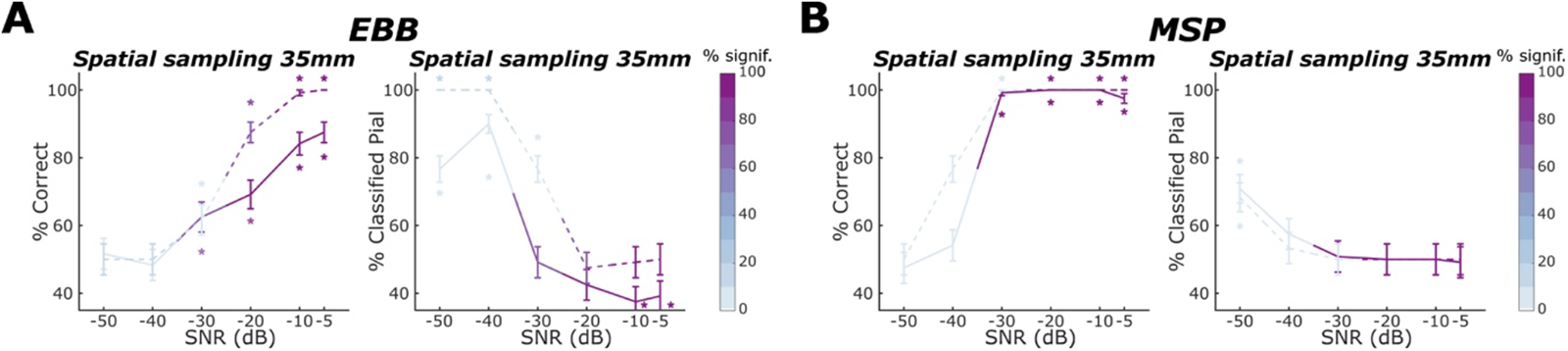
Laminar classification accuracy and bias across signal-to-noise ratios. We applied the **A** EBB and **B** MSP source reconstruction approaches to simulated data for an OPM-MEG sensor array with an inter-sensor distance of 35 mm. Solid lines denote laminar inference based on the whole-brain free energy analysis; dashed lines denote laminar inference based on the ROI-t-statistic analysis. Left columns within each subpanel show the percentage of correct laminar inferences, the right columns show the percentage of simulations where laminar inference favoured the pial source model. The percentage of simulations with free energy differences or t-statistics exceeding the significance threshold is represented by the intensity of the line colour. The error bars represent the standard error. Asterisks show where the percentage is significantly above or below chance levels. **A** For the EBB approach, we found significant increases in classification accuracy with increasing SNR. **B** For the MSP approach, we observed an excellent classification performance with accuracy at ceiling and no biases for SNRs of −30 dB or higher for both, free energy and ROI-t-statistic, analyses.

Results across OPM sensor arrays with varying inter-sensor distances can be found in the Supplementary section in Figure S1. We replicate previous results^26^ that we cannot successfully distinguish between superficial and deep sources using minimum norm and LORETA source localisation approaches (see Supplementary Fig. S2).

### Increasing sensor sampling density

We next investigated the impact of the sensor sampling density, here parameterised by the inter-sensor distance, on classification performance. Results are summarised in Figure 2. For the EBB approach combined with the whole-brain free energy analysis, classification accuracy significantly decreased with increasing inter-sensor distances at SNRs of −5 dB (beta = −0.710, p <.01), −10 dB, (beta = −0.781, p <.001) and −20 dB (beta = −0.558, p = 0.001), but not at lower SNRs where laminar inference was challenging or not feasible at all. Sampling density had no discernible impact on classification bias, except at very low SNRs, where classification bias towards the white surface increased with increasing inter-sensor distances (−40 dB, beta = −1.398, p <.001). However, single simulations were typically not significant and classification accuracy was at chance level.

For the EBB approach combined with the ROI-t-statistics analysis, classification accuracy was at ceiling across sensor sampling densities at a high SNR of −5 dB and decreased significantly with increasing inter-sensor distances at SNRs of −10 dB (beta = −1.692, p <.01), −20 dB, (beta = −1.178, p <.001) and −30 dB (beta = −0.522, p <.01).

Increasing inter-sensor distance led to a significant increase in bias towards the pial surface at −20 dB (beta = 0.356, p <.05) and −30 dB (beta = 2.080, p <.001). As described earlier, classification at the single simulation level was reliably significant only for sensor array configurations with dense spatial samplings at −20 dB and not feasible at any spatial sampling densities at an SNR of −30 dB. At an SNR of −40 dB, the ROI-based t-statistic classification was extremely biased towards the pial surface across all sensor sampling densities. While the free energy metric was biased towards the pial surface as well, this bias decreased with decreasing sensor counts, i.e., it was less pronounced at lower sampling densities.

For the MSP approach, at SNRs of −30 dB or higher, classification performance was at ceiling and showed no significant biases. Classification performance and bias did not vary significantly across sampling densities. At these moderate to high SNRs, the whole-brain and ROI-based analyses yielded similar results as indicated by non-significant exact McNemar’s tests. At −30 dB, we observed a trend towards decreased classification accuracy with increasing inter-sensor distances for the free energy analysis (logistic regression: beta = −1.547, p = 0.082), while laminar inference tended to be non-significant at the single source level for the ROI analysis.

At −40 dB, we observed a steep decline in performance accuracy with decreasing sensor counts (beta = −1.455, p <.001). However, as mentioned earlier, the corresponding classifications were not reliably significant at the level of single simulations, i.e., sources.

Overall, in our simulations, we observed significant decreases in classification accuracy as the inter-sensor distances increased for the EBB source reconstruction approach. In contrast, the MSP approach demonstrated a consistent classification performance close to ceiling across varying inter-sensor distances at SNRs of −30 dB and higher.

**Fig. 2.**
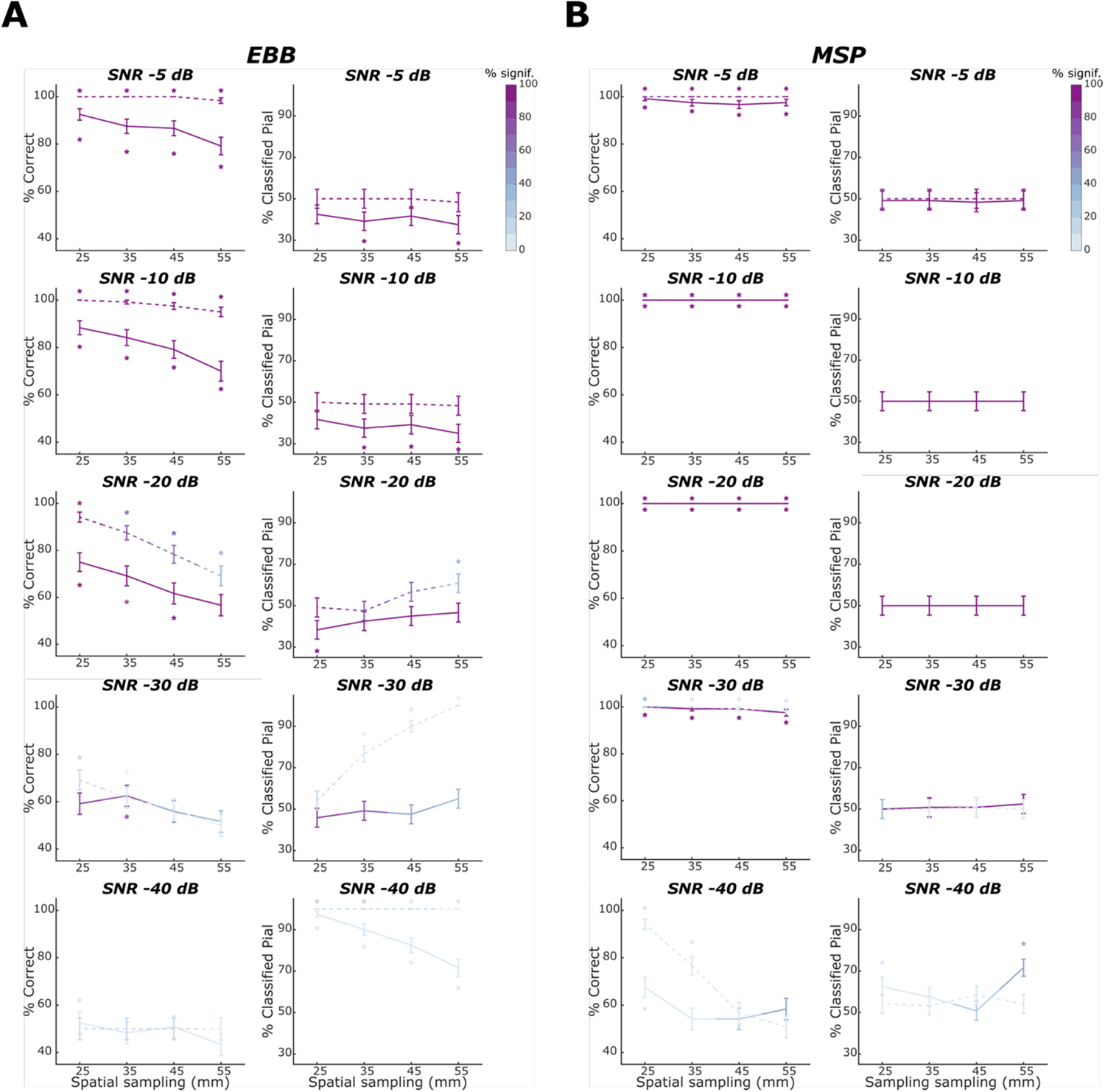
Laminar classification accuracy and bias across OPM array inter-sensor distances. Solid lines denote laminar inference based on the whole-brain free energy analysis; dashed lines denote laminar inference based on the ROI-t-statistic analysis. Left columns within each subpanel show the percentage of correct laminar inferences, the right columns show the percentage of simulations where laminar inference favoured the pial source model. SNR decreases across rows. The percentage of simulations with free energy differences or t-statistics exceeding the significance threshold is represented by the intensity of the line colour. The error bars represent the standard error. Asterisks show where the percentage is significantly above or below chance levels. **A** For the EBB approach, we found significant decreases in classification accuracy with increases in inter-sensor distances at SNRs of −20 dB or higher. **B** For the MSP approach, we observed no significant changes in laminar classification performance across sampling densities at SNRs of −30 dB or higher.

### Increasing the number of measurement axes

To investigate the impact of the number of measurement axes on laminar inference performance, we kept the sampling density fixed and varied the number of measurement axes. In Figure 3 we report results for a simulated OPM-MEG array with 55 mm inter-sensor distance, which corresponds to 29, 58 and 87 channels for single axis, dual axis and triaxial sensors, respectively. Note that the channel counts for dual and triaxial sensors deviate from being multiples of the sensor counts of the single axes due to a random factor in the point packing algorithm. The point packing algorithm was re-initialized for each array configuration, leading to the observed deviations in channel counts.

We found that classification accuracy increased significantly with the number of measurement axes at SNRs of −20 dB or higher when using the EBB approach combined with the free energy analysis (SNR −5 dB: beta = −0.578, p <.05; SNR −10 dB: beta = −0.709, p < .01, SNR −20 dB: beta = −0.500, p < .05). At these SNRs, we observed a bias towards the deep surface for the whole-brain analysis. This bias did not systematically increase or decrease with an increasing number of measurement axes. We found no advantage of increasing the number of measurement axes at SNRs of −30 dB and −40 dB. At −40 dB, we observed a bias towards the pial surface, which increased significantly with the number of measurement axes (beta = −1.767, p < .001). Note that while this bias was strong, the underlying differences in free energy were not significant at the single source level, i.e., the absolute log free energy differences did not exceed the significance threshold of 3. For the ROI-based analysis, classification accuracy was close to ceiling for an SNR of −5 dB and increased with the number of measurement axes at −10, −20 and −30 dB (−10 dB: beta = −2.579, p <.05; −20 dB: beta = −1.233, p < .001; −30 dB: beta = −0.409, p<.05). At an SNR of −30 dB, we observed a bias towards the pial surface which decreased significantly with added measurement axes (beta = 1.792, p < .001). However, laminar inference was not statistically significant at the single simulation level. At a very low SNR of −40 dB, classification accuracy was at chance level with an extreme bias towards the pial surface. Note that again these laminar inferences were not significant at the single source level.

For the MSP approach, both whole-brain and ROI-based analyses performed near-ceiling for SNRs of −30 dB or higher, irrespectively of the number of measurement axes. At an SNR of −40 dB, classification accuracy increased strongly with the number of measurement axes for the ROI analysis (ROI: beta = −1.148, p < .001); however, these laminar inferences did not exceed the significance threshold at the single simulation level.

We anticipated that the advantage of a more homogenous spatial coverage provided by sensors with multiple axes would be particularly evident for sparse OPM sensor arrays. Results indicating comparable yet less pronounced effects for a more densely arranged OPM sensor array can be found in Fig. S3.

In summary, our findings showed significant improvements in classification accuracy with an increasing number of measurement axes for the EBB source reconstruction approach. Conversely, the MSP approach exhibited classification performance close to ceiling levels for SNRs of −30 dB or higher, regardless of the number of measurement axes used.

### Impact of increasing scalp-sensor offsets

To test for the impact of sensor-scalp offsets, we simulated an OPM sensor array with 35 mm inter-sensor distances and single radial axis configuration and increased the scalp-sensor offsets from 6.5 to 20, 30 and 40 mm. We expected a higher classification accuracy at sensors closer to the scalp, as being closer to the cortical surface (1) renders the ratio of the distances of the sensors to the deep and superficial surfaces larger and (2) results in larger lead fields, which have been linked to improved laminar inference performance^26^.

For the EBB approach (Fig. 4A) combined with the free energy analysis, increasing the scalp-sensor offset at a high SNR of −5 dB yielded a higher classification accuracy and a weaker bias towards the deep surface. While these effects were not statistically significant (accuracy: beta = −0.423, p = 0.130; bias: beta = −0.190, p = 0.249), this was a surprising finding given the expected benefit of moving the sensors closer to the scalp and thus the cortical sources. At an SNR of −10 dB, using the same approach, classification accuracy did not change markedly across offsets (beta = 0.534, p = 0.472), while at −20 and −30 dB classification accuracy decreased with increasing scalp-sensor offsets. Again, these decreases were not statistically significant (SNR −20 dB: beta = 0.171, p = 0.320.; SNR −30 dB: beta = 0.165, p = 0.320).

For the ROI-based classification metric, classification accuracy was at ceiling at high SNRs of −5 and −10 dB and decreased significantly with increasing scalp-sensor offset at −20 dB (beta = 0.467, p = .032). At an SNR of −30 dB, classification accuracy was generally poor, ranging from 56.67 to 62.50%, and did not vary significantly across scalp-sensor offsets, while the bias towards the pial surface decreased significantly with scalp-sensor offset distance (beta = 0.613, p < .001). However, laminar inference was not significant at this low SNR, i.e., single source classifications did typically not exceed the significance threshold. At an SNR of −40 dB, classification was at chance level with a strong bias towards the pial surface for both analyses, irrespective of the scalp-sensor offsets. Again, laminar inferences were not significant at the single simulation level.

For the MSP source reconstruction approach (Fig. 4B), classification accuracy performed at ceiling for SNRs of −30 dB or higher for both, the whole-brain and the ROI-based analyses, and thus did not vary significantly across scalp-sensor offsets. At an SNR of −40 dB, we observed a significant increase towards a bias to the pial surface with increasing scalp-sensor offsets (whole-brain: beta = −0.490, p <.01; ROI: beta = −0.346, p = 0.038). However, laminar inferences at −40 dB were not significant at the level of single simulations irrespective of the classification metric or the scalp-sensor offset.

**Fig. 3.**
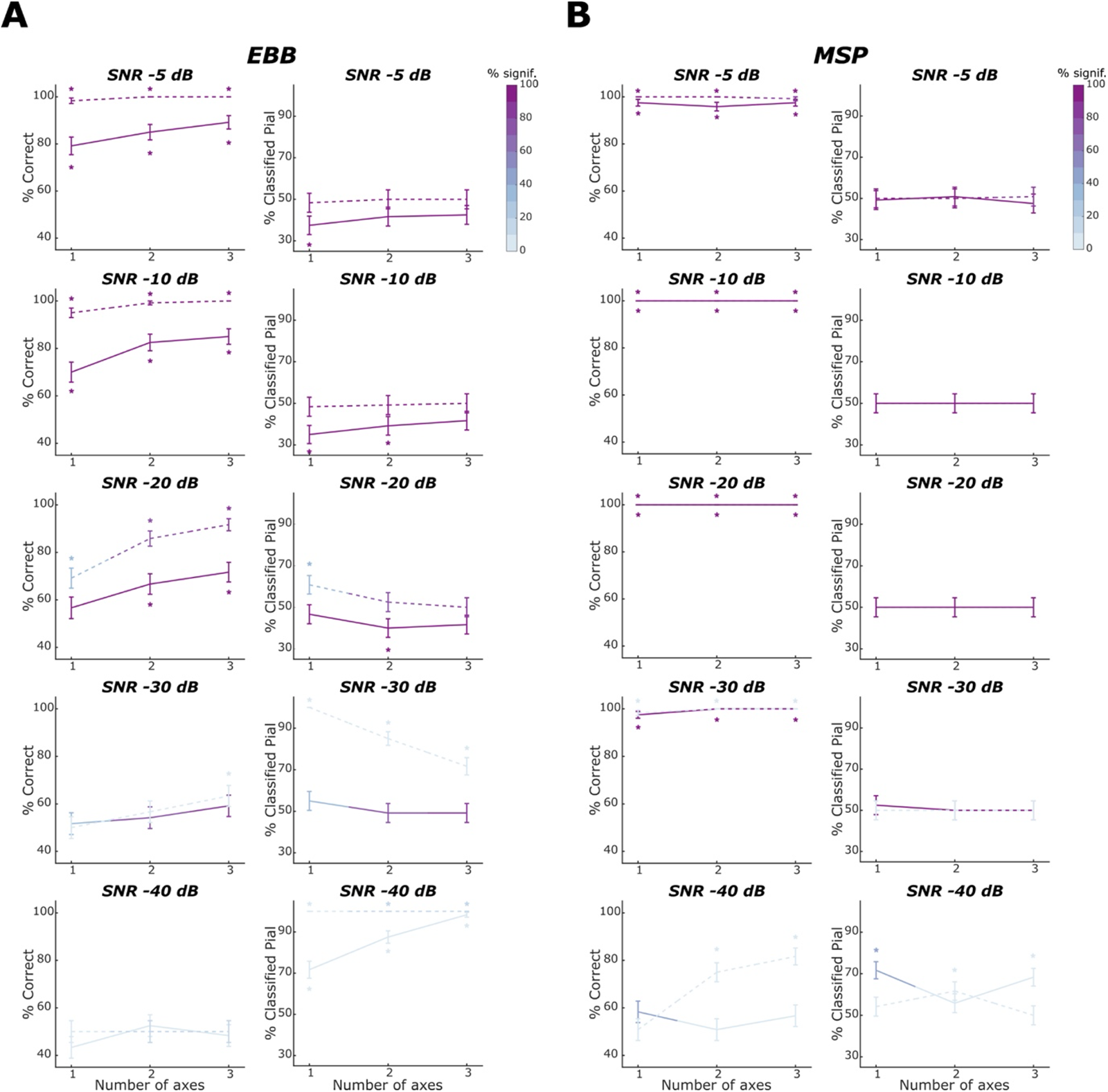
Laminar classification accuracy and bias across number of measurement axes for an OPM-MEG array with a sensor spacing of 55 mm. Solid lines denote laminar inference based on the whole-brain free energy analysis, dashed lines laminar inference based on the ROI-t-statistic analysis. Left columns within each subpanel show the percentage of correct laminar inferences, right columns show the percentage of simulations where laminar inference favoured the pial source model. **A** For the EBB approach, we found significant increases in classification accuracy with increases in the number of measurement axes at SNRs of −30 dB or higher. **B** For the MSP approach, both free energy and ROI-based analyses performed at ceiling for SNRs of −30 dB or higher, irrespectively of the number of measurement axes.

Our finding, although not statistically significant, that classification accuracy improved with increasing sensor-scalp offsets at a high SNR of −5 dB when using the EBB approach, was surprising, given the expected advantage of moving the sensors closer to the scalp. We speculate that the increase in classification accuracy with greater scalp-sensor distances at high SNRs might be due to the reduced aliasing of higher spatial frequency information when sensors are positioned farther away from the brain sources. Since sparser sensor arrays are more susceptible to aliasing, we expect this effect to be more pronounced in sparse arrays and less pronounced in dense sensor arrays. Simulations involving OPM-MEG arrays with inter-sensor distances of 55 mm and 25 mm partially support this hypothesis (Fig. S4).

**Fig. 4.**
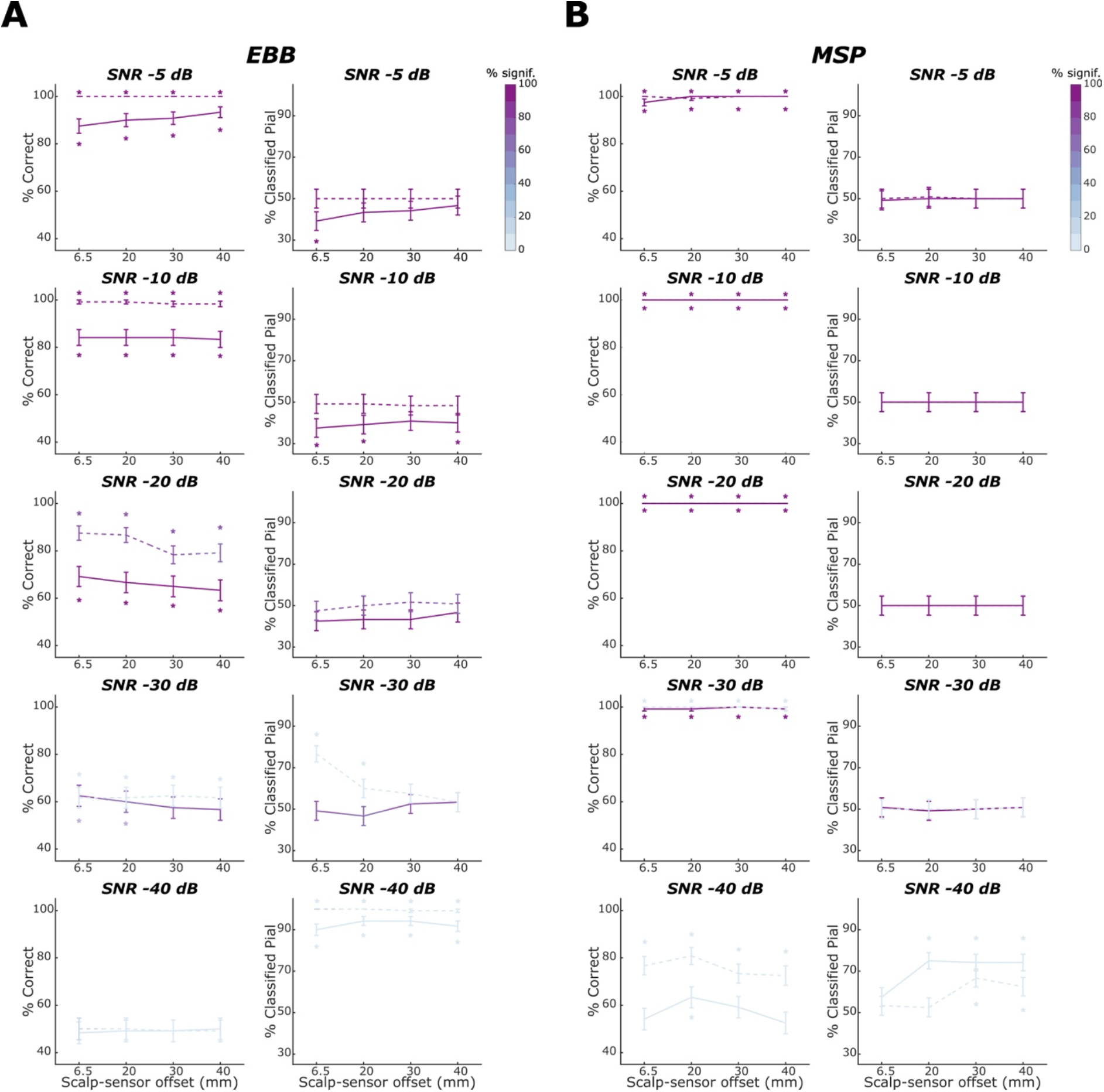
Laminar classification accuracy and bias across scalp-sensor offsets for an OPM-MEG array with a sensor spacing of 35 mm. Solid lines denote laminar inference based on the whole-brain free energy analysis; dashed lines laminar inference based on the ROI-t-statistic analysis. Left columns within each subpanel show the percentage of correct laminar inferences, right columns show the percentage of simulations where laminar inference favoured the pial source model. **A** For the EBB approach, we observed a tendency of increases in scalp-sensor distances leading to an increasing laminar classification accuracy at high SNRs and a decreasing one at a lower SNR of −20 dB. **B** For the MSP approach, classification accuracy performed at ceiling for SNRs of −30 dB or higher for the whole-brain and the ROI-based analyses and did not vary significantly across scalp-sensor offsets.

### Co-registration errors

To investigate the impact of co-registration errors on our ability to perform non-invasive laminar inference, we ran simulations with a single-axis array at an inter-sensor distance of 35 mm and an SNR of −10 dB and added small random displacements to the three fiducial locations (Fig. 5). We observed a steep decrease in classification accuracy with increasing co-registration errors for the EBB approach (whole-brain: beta = 0.838, p <.001; ROI: beta = 1.189, p <.001), where laminar inference was not feasible anymore at a co-registration error of 4 mm. Classification bias remained relatively stable across co-registration errors, with a trend towards an increased bias towards the deep surface with increasing co-registration errors for the ROI-based t-statistics analysis (beta = 0.268, p = .065). For the free energy analysis, the previously observed bias towards the deep surface was sustained across co-registration errors.

For the MSP approach, classification accuracy remained high (above 90%) across co-registration errors but decreased significantly with increasing co-registration errors (whole-brain: beta = 2.497, p <.001; ROI: beta = 1.908, p <.01). No classification biases were observed for either the whole-brain or the ROI-based analysis.

**Fig. 5.**
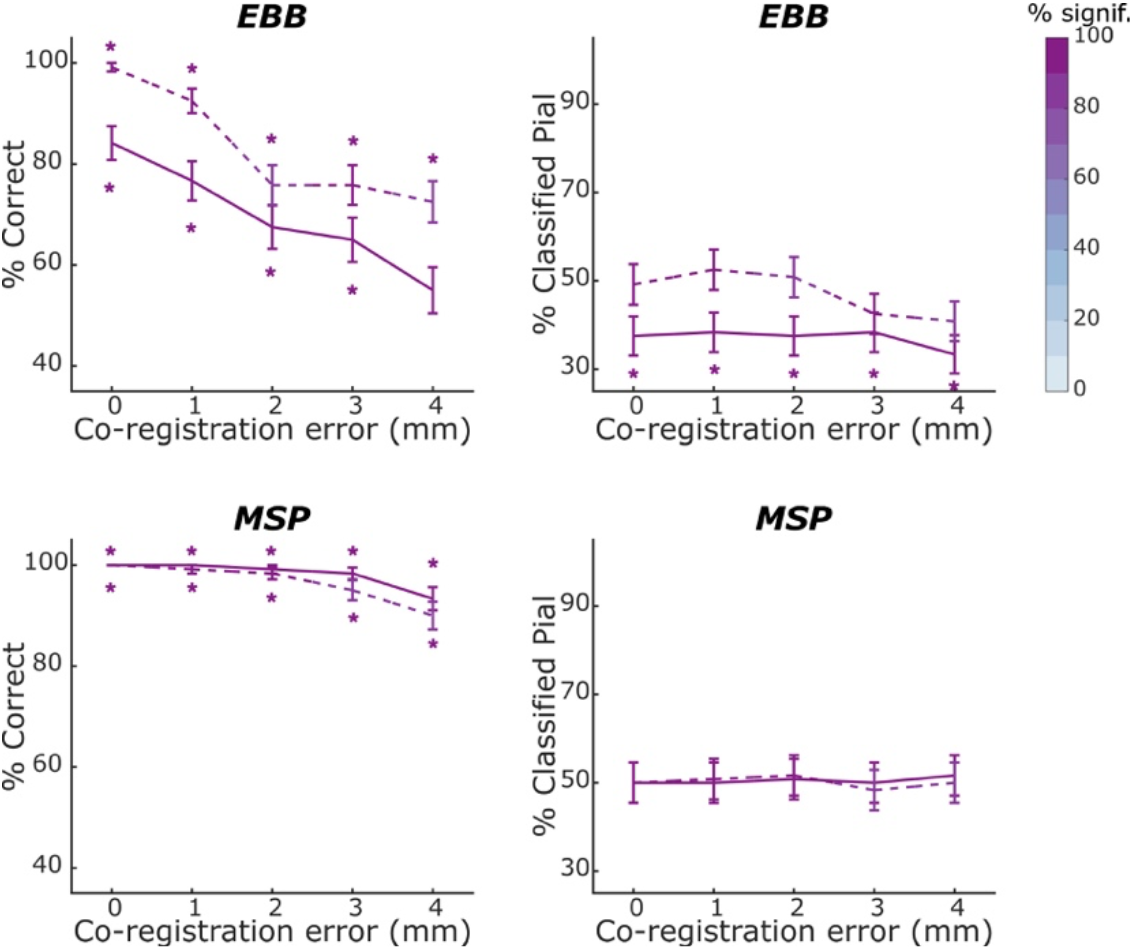
Laminar classification accuracy and bias across co-registration errors at an SNR of −10 dB. We used an OPM-MEG array with a sensor spacing of 35 mm and applied the EBB and MSP source reconstruction approaches to the simulated data. Solid lines denote laminar inference based on the whole-brain free energy analysis; dashed lines laminar inference based on the ROI-t-statistic analysis. Left columns show the percentage of correct laminar inferences, right columns show the percentage of simulations where laminar inference favoured the pial source model. For the EBB approach, we observed a decrease in classification accuracy with increasing co-registration errors. We did not observe any significant changes in classification bias with varying co-registration errors for either the whole-brain or the ROI-based analysis. For the MSP approach, classification accuracy decreased significantly with increasing co-registration errors. No classification biases were observed for either the whole-brain or ROI-based analyses.

### Incongruent patch sizes

To test for the impact of incongruencies between the true and the assumed source extent, we simulated congruent and incongruent source and reconstruction patch sizes of 5 and 10 mm for a single-axis OPM-sensor array with an inter-sensor distance of 35 mm and performed laminar inference using the whole-brain free energy analysis.

For the EBB source reconstruction approach, we found a significant drop in laminar classification accuracy for over- (at SNRs of −5, −10 and −20 dB: p < .01, p < .01 and p < .05, respectively) and underestimated (at SNRs of −5 and − 10 dB: both p < .00001) patch sizes (Fig. 6). At SNRs of −5 dB and −10 dB, simulations with congruent patch sizes (shown in purple) were biased towards deep layers as observed in our previous simulations (Fig. 1-4). Compared to congruent patch sizes, overestimation of patch sizes yielded a shift towards pial classification (rendering the classification bias non-significant), while underestimation of patch sizes led to an even stronger bias towards the deep cortical surface. However, two-sided exact McNemar’s tests indicated that these shifts were not statistically significant at any of the SNRs tested.

For the MSP approach, we found smaller but significant decreases in classification accuracy for incongruent patch sizes (over-estimated patch sizes at SNRs −10, −20 and −30 dB: p <.05, p <.01 and p <.05; under-estimated patch sizes at SNRs of −20 and −30 dB: p < .05 and p < .001, respectively) and no classification bias for neither congruent or incongruent patch sizes at all SNRs that enable statistically significant laminar inference (−30 dB or higher).

**Fig. 6.**
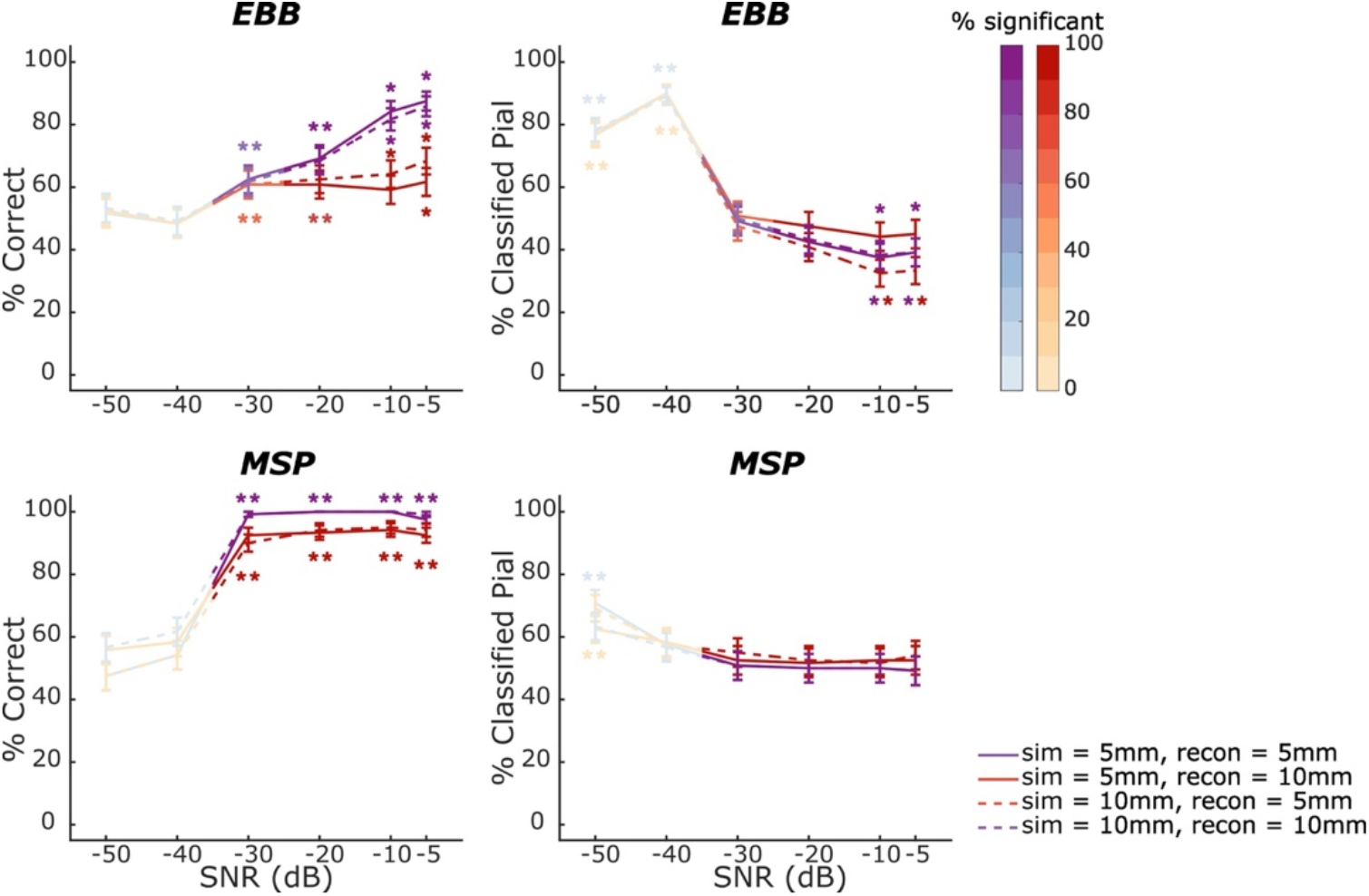
Whole-brain free energy analysis laminar inference with congruent and incongruent patch sizes for an OPM-MEG array with a sensor spacing of 35 mm. Purple lines denote simulations where the reconstructed patch size matches the simulated patch size (solid = 5 mm, dashed = 10 mm), red lines are where patch size is either over-(solid) or underestimated (dashed). The percentage of simulations with free energy differences or t-statistics exceeding the significance threshold is represented by the intensity of the line colour. Asterisks show where the percentage is significantly above or below chance levels. The left column shows the percentage of sources classified correctly for each level of SNR tested. The error bars represent the standard error. As SNR increased, classification accuracy was reduced for both source reconstruction approaches when the patch size was under- or overestimated. The right column shows the percentage of sources classified as coming from the pial surface for each level of SNR tested. As SNR increased, congruent patch sizes resulted in a bias towards the deep surface for the EBB approach. Overestimation of patch sizes reduced this bias, while underestimation of patch sizes led to an even stronger bias towards the deep surface. The MSP approach showed no bias as SNR increased, irrespective of whether congruent or incongruent patch sizes were used.

### Interfering brain noise sources

Next, we investigated the impact of internal noise (nuisance) sources on our ability to accurately infer the laminar origin of the simulated sources. To this end, we modelled sensor activity from a laminar cortical source of interest at an SNR of −5 dB and added concurrent weaker noise sources on the mid cortical surface. We simulated the sensor activity for a dense OPM-MEG array with an inter-sensor distance of 25 mm and single axis sensors and again applied the whole-brain and ROI-based laminar inference analyses for EBB and MSP source reconstruction approaches. Results are summarised in Fig. 7.

For the EBB source reconstruction approach and the whole-brain analysis, we were not able to infer the laminar origin of the simulated sources (correct = 54.17%, p = ns.) and observed a strong bias to the deep surface (white matter = 87.50%, p <.00001). This contrasts with our results for the ROI analysis, which showed a high classification accuracy (EBB: correct = 91.67%, p <.00001) and no bias (pial = 50.00%, ns.). For the MSP source reconstruction approach, the whole-brain analysis similarly showed a low classification accuracy, which however was still above chance level (correct = 60.83%, p <.05), and laminar inference was biased towards the deep surface (white matter = 76.67%, p <.0001). For the ROI analysis, the MSP approach yielded a high classification accuracy (correct = 93.33%, p <.00001) and showed no classification bias (pial = 51.67%, ns.).

**Fig. 7.**
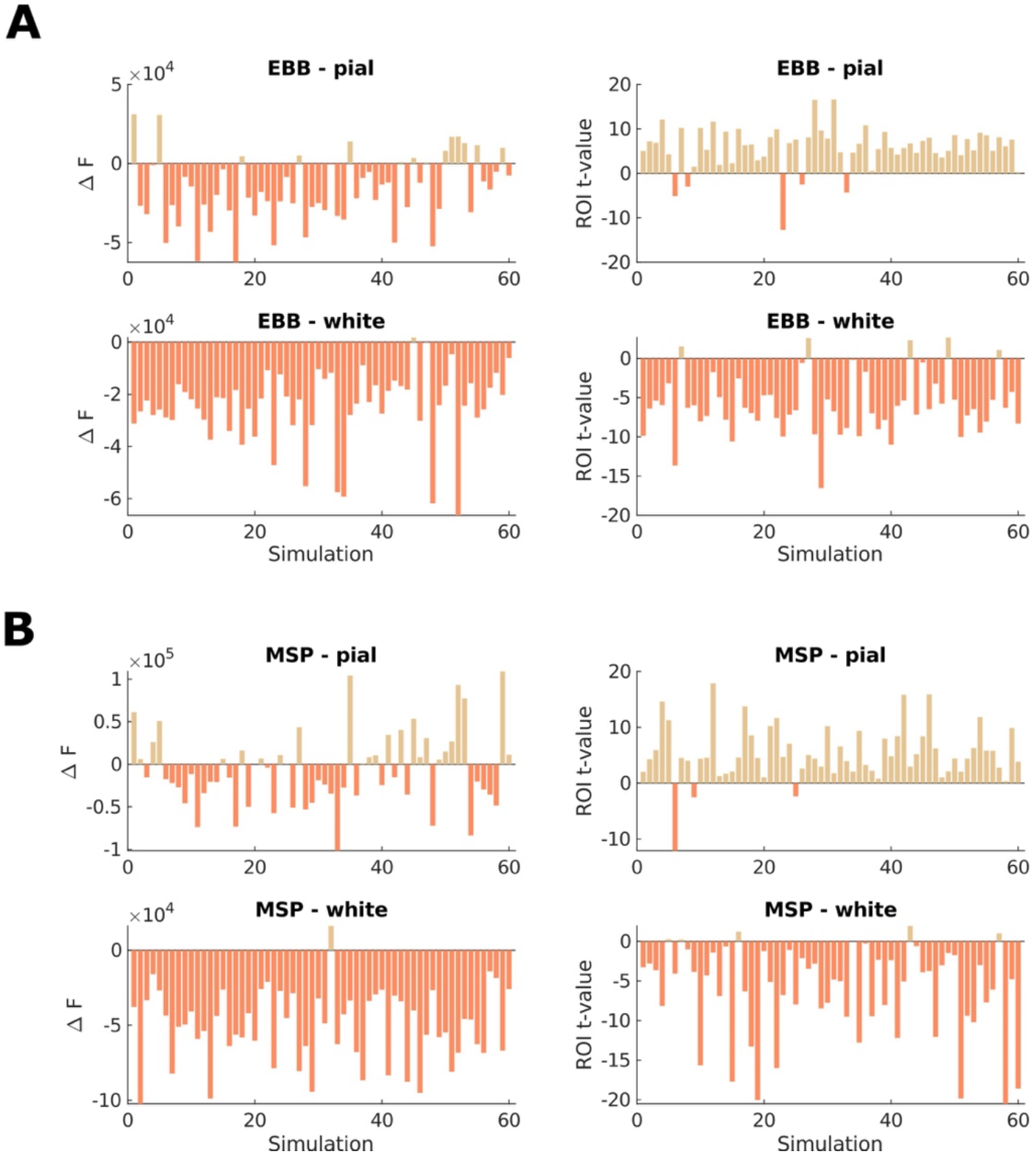
Laminar source discrimination in the presence of interfering brain noise sources. We used a dense OPM-MEG array with an inter-sensor distance of 25 mm and simulated sensor activity at a relatively high SNR of −5 dB to mimic a best-case scenario. Left column: The difference in free energy between the pial and white matter generative models in each simulation. Right column: t-values for the difference between pial and white matter ROI values for each simulation. Each panel shows simulations with pial surface sources on the top row, and simulations with white matter surface sources on the bottom row. For the **A** EBB and **B** MSP source reconstruction approach, the whole-brain analysis similarly showed a low classification accuracy and a bias towards the deep surface (white matter = 76.67%, p < .0001), while the ROI-based analyses yielded high classification accuracies and showed no classification bias.

## Discussion

We evaluated the efficacy of high-accuracy forward models in achieving laminar discrimination using on-scalp OPM sensors. Remarkably, we found that adequate laminar inference can be achieved with relatively few sensors at coarse spatial samplings (> 50 mm) and moderate to high SNRs. For the EBB approach, classification accuracy improved with increasing spatial sampling densities and measurement axes. For the MSP approach, classification performance was near ceiling for SNRs of −30dB or higher and unattainable at lower SNRs, regardless of the spatial sampling employed. We note that the excellent performance of the MSP approach at SNRs above −30dB can be partially attributed to the distinct advantage of its source space priors, which include the patches where sources were simulated.

As expected, we observed that laminar inference was vulnerable to over- and underestimation of source patch sizes, co-registration errors and interfering brain sources. Interestingly, contrary to our expectations, moving the sensors closer to the scalp, which is arguably the game-changing feature of room-temperature OPM sensors, did not provide as much benefit for successful laminar inference as expected and even had a harmful effect at high SNRs. The biases observed in laminar inference also warrant caution. We found strong biases towards both the superficial and deep surfaces, particularly at very low SNRs, although these biases did not typically reach statistical significance. Furthermore, the EBB approach, when combined with whole-head free energy analysis, exhibited a notable bias towards the deep surface, particularly at high SNRs. Since at high SNRs, beamformer images tend to be more focal^50^, we propose that the denser sampling of the deep surface could be advantageous in such scenarios.

Based on our findings, we recommend using either the MSP approach or adopting the ROI-based analysis for the EBB approach, as it demonstrated higher classification accuracy than the whole brain analysis without exhibiting a bias towards the white surface. Regardless of the chosen approach, ensuring adequate SNR through an appropriate number of trials and shielding, as well as maintaining precise co-registration accuracy is crucial. Bayesian model comparison across various patch sizes can assist in the decision-making process of selecting a suitable patch size^27^. In the following section, we will discuss the results in more detail.

### Number of sensors

As expected, increasing spatial sampling density led to increased classification accuracy, in line with previous studies that have shown increased spatial resolution for source-reconstructed activity with increasing sensor counts for SQUID-MEG^30^ and OPM-MEG^7,9,31^. However, our finding that laminar inference is possible at a low sensor count of 32 sensors was somewhat surprising given that previous studies have suggested that approximately 300 sensors are needed to achieve a spatial discrimination of 2 to 2.5 mm^9,40^. While even this spatial resolution is below what is assumed to be required to distinguish between deep and superficial current dipoles, we argue that employing high-accuracy laminar forward models helped to boost the effective spatial resolution.

Unlike the whole-brain analysis, the ROI-based analysis could still recover the simulated signal at −30dB, however with decreasing accuracy and increasing bias towards the pial surface as the inter-sensor density increased. As we used uncorrelated sensor noise in our simulations, the sensor noise has a high spatial frequency. It is thus more easily fitted using the pial surface as this surface tends to be closer to the sensors and pial sources are expected to yield more focal sensor patterns. At sparser sensor samplings the simulated dipole current sources cannot be recovered, and the signal is dominated by sensor noise, which could explain the increase in bias towards the pial surface with increasing inter-sensor distances. For the whole-brain analysis and an SNR of -40dB, the simulated dipole current sources cannot be recovered at any of the sensor samplings tested and the decrease in bias towards the pial surface may be due to the fact the lower spatial frequency of the sensor noise pattern for sparse sensor arrays can be more easily explained by sources on the deep surface as the sensor noise patterns for dense arrays. It is important to note that while these biases were pronounced, classification at the single simulation level did in general not reach significance at such low SNRs.

### Number of measurement axes

Classification accuracy increased with the number of measurement axes, consistent with findings that increasing the number of axes results in increased information content^10^ and better spatial coverage^32,33^. As expected, the benefit of multiple axes was more pronounced at lower inter-sensor distances, where undersampling of high spatial frequency features leads to unexplained noise due to signal aliasing. Note that our simulations did not include external noise sources, and the advantage of dual-axis and triaxial sensors is likely to be even greater in the presence of such interference^32^.

### Impact of sensor-scalp offset distance

One of the fundamental assumptions behind the postulated potential of OPMs for non-invasive electrophysiology is that sensors closer to the scalp will increase the sensitivity to cortical sources, resulting in higher information capacities and spatial resolution as well as better source separability compared to conventional MEG^10,40,51^. However, our simulations indicate that at high SNRs increasing the scalp-sensor offset increases classification accuracy when using the EBB approach combined with the free energy analysis.

We set out to test the hypothesis that this increase in laminar inference accuracy at larger scalp-sensor offsets could be attributed to stronger aliasing effects for on-scalp sensor arrays. Aliasing due to undersampling limits the effective SNR of on-scalp sensors^40^. For a lower number of sensors, where aliasing of higher spatial frequencies is more pronounced, it may thus be beneficial to increase the distance from the scalp to achieve better spatial coverage. We thus predicted an even stronger advantage of large scalp-sensor offsets for sparser arrays and no advantage for denser arrays. While our predictions were confirmed for the sparser array, the benefit of larger scalp-sensor offsets remained present for the denser array (See Fig. S4 in “Supplementary material”). Therefore, aliasing effects are unlikely to fully account for our findings.

What explains the better laminar inference at larger offsets at high SNRs and how do we account for the reversal of scalp-sensor offset effects with decreasing SNRs? While a high SNR corresponds to low noise interference, for sparse on-scalp sensor arrays the actual signal may not be adequately sampled and we may thus benefit from moving the sensors further away from the scalp. This reduces aliasing, which increases the effective SNR and hence classification accuracy. At lower SNRs, we must deal with stronger sensor noise and may struggle to pick up the simulated current sources at all if we move the sensors too far away from the scalp. This complex interplay between SNR, sampling sensor densities, and scalp-sensor offsets requires further investigation. However, we emphasise that OPM sensor arrays, unlike SQUID sensor arrays, at least allow for the adjustment of scalp-sensor distances. We further note that here we compared laminar classification performance across scalp-sensor offsets at fixed SNRs and did not account for any expected increases in SNR when moving sensors closer to the scalp.

### Impact of co-registration errors

Both source reconstruction approaches showed declines in classification accuracy with increasing co-registration errors. However, laminar inference remained possible for co-registration errors of up to 4 mm, depending on SNR, sensor density, as well as the source reconstruction approach and laminar inference analysis used.

Here we modelled co-registration errors for a rigid sensor array, i.e., we assume systematic shifts due to fiducial localisation errors like in SQUID-MEG, rather than random sensor localisation or orientation errors. The latter can have a more detrimental effect on source reconstruction accuracy^14^. Zetter et al. (2018)^41^, using random displacement errors, proposed a cut-off of 4 mm sensor position and 10° sensor orientation RMS errors for acceptable mis-co-registration, noting that at larger co-registration errors, the advantage of on-scalp MEG may be lost. Troebinger et al. (2014)^23^, in a simulation study based on a SQUID-MEG rigid helmet, added rotation or pure translation to fiducial locations and suggested a cut-off of less than 2 mm/2° of co-registration error for accurate laminar inferences. Localising sensors with this degree of accuracy is challenging but feasible, as shown for co-registration between on-scalp MEG and MRI images^42,52^. Even higher accuracy, with errors below 0.5 mm, may be necessary for localising multiple, dependent sources with added noise sources^7^. For experimental setups, the use of rigid measurement helmets in conjunction with co-registration devices with an excellent accuracy, such as structured-light or laser scanners, will be critical to reach the necessary co-registration accuracies.

Co-registration errors are not the only errors that will affect forward model accuracy and, consequently, localisation accuracy. It is worth noting, that we did not evaluate the impact of crosstalk and gain errors (Duque-Munoz et al., 2019^53^), orientation errors, or cross-axis projection errors^13,54,55^.

### Patch size incongruencies

We have replicated previous findings that over- or under-estimation of patch sizes can decrease classification accuracy and bias laminar results^23,25^. Compared to congruent patch sizes, overestimation of patch sizes resulted in a shift towards pial classification, rendering the classification bias non-significant, while underestimation of patch sizes led to an even stronger bias towards the deep surface. We note that these shifts in bias with incongruent patch sizes were not statistically significant at any SNR as evaluated with the exact McNemar’s test. However, these results are consistent with earlier findings, and we interpret them as follows: larger patch sizes and deeper sources both lead to spatially more spread-out sensor signals, and underestimation of patch sizes and the resulting unexpected lower spatial frequencies at the sensor level can be explained by assigning a source to the deep cortical surface. Similarly, a sensor topography more focal than expected from overestimated patch sizes can be explained by assigning a source to the superficial surface^23^. Even when the ground truth on the source extent is unknown for experimental data, the optimal patch size for the laminar inference procedure can be determined by comparing the model evidence of combined pial/white matter source inversions across varying reconstruction patch sizes^27^.

### Interfering brain noise sources

We next investigated laminar classification performance in the presence of five interfering noise sources on the mid cortical surface. We observed that, despite employing a favourable setup with high SNR and dense sensor sampling, the whole-brain analysis was unable to successfully recover the laminar origin of the source of interest. Performance accuracy of the ROI-based approach remained high and did not show any bias in the presence of noise sources. We note that the laminar source of interest had a larger SNR than the interfering noise sources, and the activity-based ROI was thus mostly defined by the source of interest, effectively masking the noise sources. We further highlight that while we modelled the interfering brain noise sources at the mid cortical layer, no surface for the mid cortical layer was incorporated into the high-precision forward model. Such forward models with a more fine-grained laminar resolution offer an exciting perspective for future studies.

### Alternative approaches to infer laminar MEG activity

We were able to successfully infer the laminar origin of simulated current dipoles using OPM-MEG sensor arrays at comparatively low spatial sampling densities. Nevertheless, in the light of our complex results regarding the impact of varying scalp-sensor offsets, and the observed classification biases, the potential of OPM-MEG to improve the accuracy of laminar inferences may be less striking than initially presumed. This is at least the case for the laminar forward model approach investigated in the current study.

Alternative approaches to laminar inferences have been discussed previously. Pinotsis et al. (2017)^56^ and Pinotsis & Miller (2020)^57^ employed dynamical causal modelling (DCM), with modelling parameters set based on estimates from intracranial data, and statistical decision theory to infer the laminar sources of non-invasive electrophysiological signals. However, the authors note that applying laminar DCM to non-invasive data is challenging due to the high collinearity of these parameters, and their results could not be replicated across data sets^57^. In a proof-of-principle study, Ihle et al. (2020)^29^ combined DCM and high-precision forward models to recover the laminar origin of a cortical current source. Yet, their approach was restricted to a single dipole pair at a known spatial location, limiting their applicability to more realistic scenarios.

A promising path towards non-invasive laminar inference could be to combine classical dipole fitting or beamformer source reconstruction with high-density OPM arrays. Recent simulation work^7^ estimated that a densely packed magnetocorticography array of 56 OPM sensors would be able to localise multiple electrophysiological brain responses at a millimetre resolution.

## Conclusion

We conducted simulations to investigate the potential of on-scalp OPM sensors combined with high-precision forward modelling for inferring the laminar origins of neural activity. Our findings provide guidance on the requirements for OPM-MEG systems in terms of sensor numbers and measurement axes to achieve robust laminar inference.

## Acknowledgements

The author is grateful to James Bonaiuto, George O’Neill, Tim Tierney and David Poeppel for useful discussions, support on implementational matters and valuable feedback on the manuscript.

## Supplementary

### Laminar inference across SNRs at varying inter-sensor distances

**Fig. S1.**
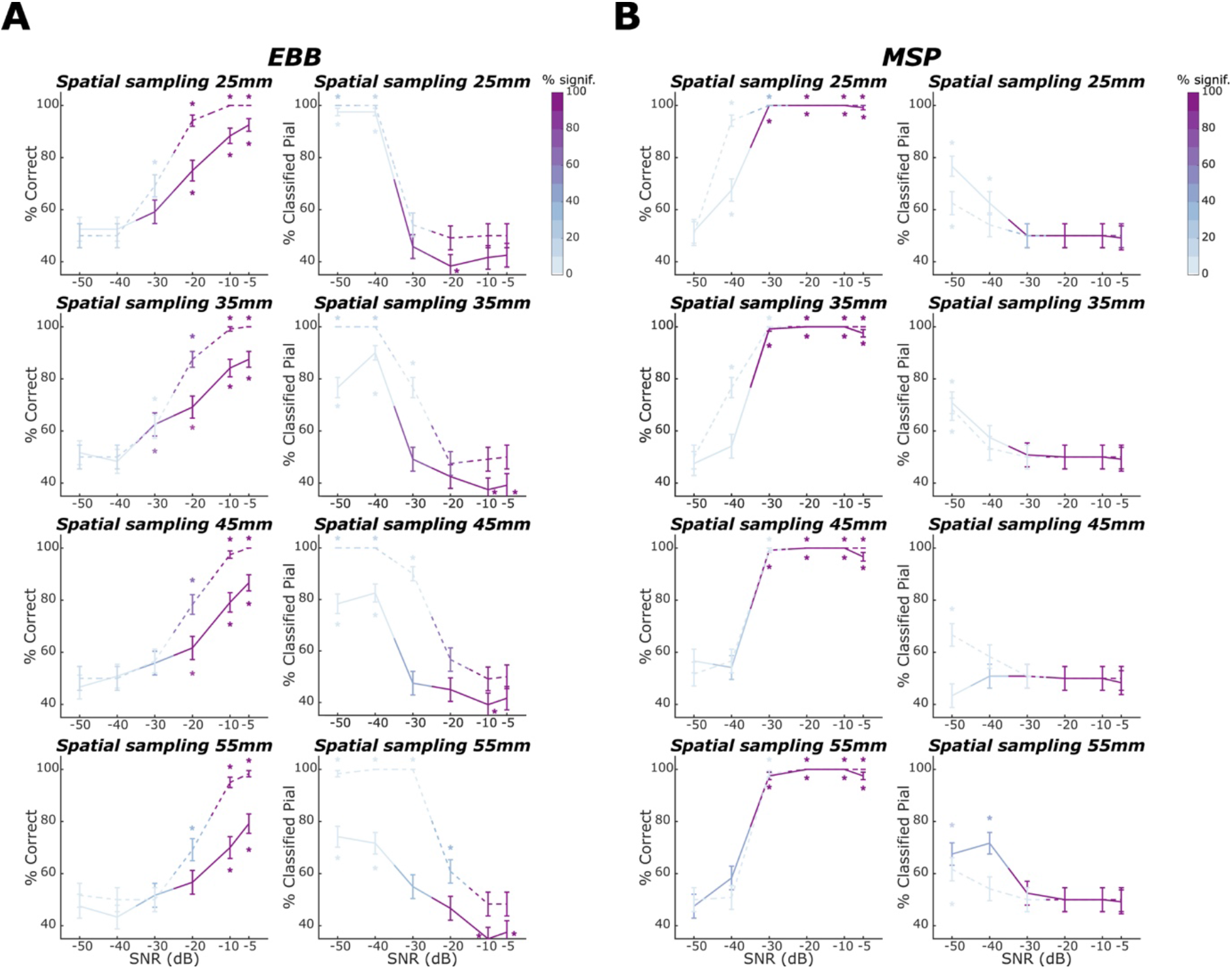
Laminar classification accuracy and bias across signal-to-noise ratios across OPM-MEG sensor arrays with varying inter-sensor distances. Solid lines denote laminar inference based on the whole-brain free energy analysis; dashed lines denote laminar inference based on the ROI-t-statistic analysis. Left columns within each subpanel show the percentage of correct laminar inferences, the right columns show the percentage of simulations where laminar inference favoured the pial source model. Inter-sensor distance decreases across rows. The percentage of simulations with free energy differences or t-statistics exceeding the significance threshold is represented by the intensity of the line colour. The error bars represent the standard error. Asterisks show where the percentage is significantly above or below chance levels. **A** For the EBB approach, we found significant increases in classification accuracy with increase in SNR for all spatial sampling distances. **B** For the MSP approach, we observed an excellent classification performance with accuracy at ceiling and no biases for SNRs of −30 dB or higher for both, free energy and ROI- t-statistic analyses.

### Laminar inference with Minimum Norm Estimates and LORETA source reconstruction approaches

We replicate previous findings that source reconstruction approaches without sparsity constraints, like Minimum Norm Estimates and Loreta, were not able to recover the laminar origin of the simulated sensor data (Fig. S2). Even under highly advantageous conditions, for a dense OPM-MEG array with an inter-sensor distance of 25 mm and simulated sensor activity at a high SNR of −5 dB, the minimum norm and LORETA source reconstruction algorithms were not able to correctly infer the laminar origin of simulated source activity using neither the whole-brain analysis (IID: correct = 54.17%, p = ns.; COH: correct = 54.17%, p = ns.) nor the ROI analysis (IID: correct = 50.00%, p = ns.; COH: correct = 50.00%, p = ns.). The whole-brain analysis was biased towards the deep surface (IID: white matter = 79.17%, p = 0.0001; COH: white = 80.83%, p = 0.0001), and the ROI-based analysis strongly towards the superficial surface (IID: pial = 100%, p < .0001; COH: pial = 100%, p <.0001). The directions of these biases replicate the findings in Bonaiuto et al., 2018.

**Fig. S2.**
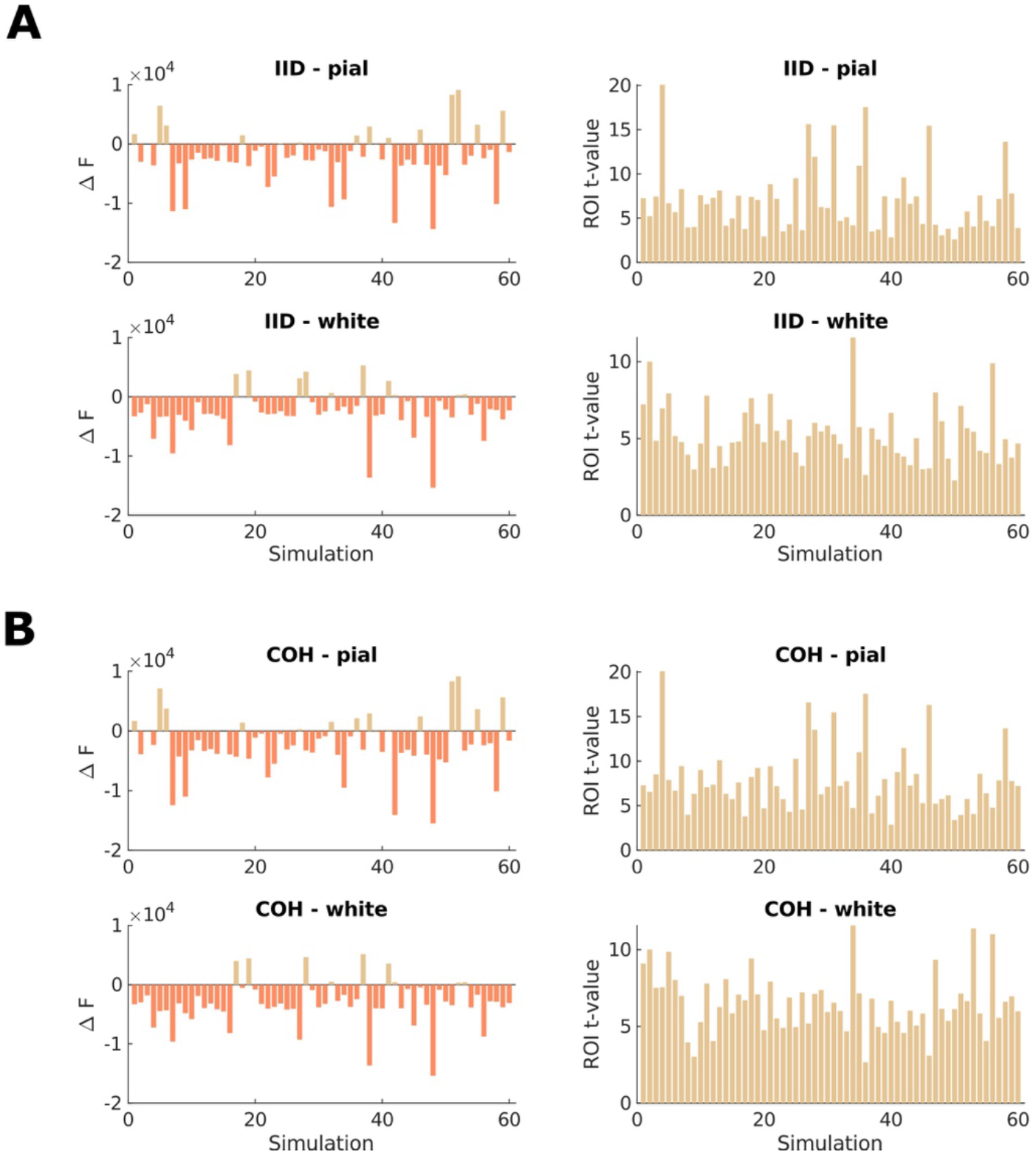
The source reconstruction approaches without sparsity constraints, i.e., MNE (IID) and Loreta (COH), were not able to perform laminar source discrimination. We used a dense OPM-MEG array with an inter-sensor distance of 25 mm and simulated sensor activity at a relatively high SNR of −5 dB. Left column: The difference in free energy between the pial and white matter generative models in each simulation (SNR = −5 dB). Right column: T-statistics from the ROI analysis comparing pial and white matter ROIs for each simulation (SNR = −5 dB). Each panel shows simulations with pial surface sources on the top row, and simulations with white matter surface sources on the bottom row. The minimum norm (**A**; IID) and LORETA **(B**; COH) source reconstruction algorithms were not able to correctly classify simulated source activity as originating from either the deep or superficial surface for the whole-brain or the ROI analysis. The whole-brain analysis was biased towards the deep and the ROI analysis strongly towards the superficial surface.

### Increasing the number of measurement axes for OPM-MEG arrays with denser spatial sampling

While we observed significant increases in classification accuracy with increasing number of measurement axes for an OPM-MEG sensor array with an inter-sensor distance of 55 mm, we acknowledge that such an effect may have been particularly strong at low sensor sampling densities where the advantage of a more homogenous spatial coverage afforded by sensors with multiple axes is expected be more pronounced. We thus re-analysed the impact of the number of measurement axes on our ability to perform laminar inference with a denser OPM-MEG array with a 35 mm inter-sensor distance and report results for 69-, 134- and 207-channel configurations for single-axis, dual-axis and triaxial sensor arrays, respectively.

For the EBB approach combined with the free energy analysis, classification performance increased with the number of measurement axes at SNRs of −20 dB or higher. However, these increases were not significant, and only trends of increasing classification accuracy were observed at −5 dB (beta = −0.603, p = 0.078) and −20 dB (beta = −0.389, p = 0.080). We again found a bias towards the deep surface, as described previously for the single-axis configuration. This bias did not systematically increase or decrease with an increase in the number of measurement axes. We found no advantage of increasing the number of measurement axes at −30 dB SNR and no significant changes in classification bias across measurement axes. At a very low SNR of −40 dB, laminar inference was not feasible, regardless of the number of measurement axes.

For the ROI-based analysis, classification performed at ceiling for SNRs of −5 and −10 dB and increased significantly with the number of measurement axes at −20 dB (beta = −1.086, p <.05). At an SNR of −30 dB, we observed a trend of increased classification accuracy with an increase in the number of axes (beta = −0.409, p = 0.051), while the bias towards the pial surface reduced with an increase in the number of measurement axes (beta = 0.641, p < .01). However, laminar inference was not statistically significant at the single simulation level. At a very low SNR of −40 dB, classification accuracy was at chance level with a bias towards the pial surface, which increased with the number of measurement axes (beta = −1.462, p < .01). Note that laminar inferences were not significant at the single source level.

For the MSP approach, both free energy and ROI-based analysis performed at ceiling for SNRs of −30 dB or higher, irrespective of the number of measurement axes. In contrast, at an SNR of −40 dB, classification accuracy increased strongly with the number of measurement axes (whole-brain: beta = −0.644, p < .01, ROI: beta = −2.186, p < .001); however, these laminar inferences did not exceed the significance threshold at the single source level.

### Impact of increasing scalp-sensor offsets for varying inter-sensor distances

We first repeated our simulations for a sparser OPM-MEG array with an inter-sensor distance of 55 mm. We expected that the improvement in classification accuracy, observed with increasing scalp-sensor distances at an SNR of −5 dB for an array with an inter-sensor distance of 35 mm, would be more pronounced. Additionally, we expected the benefit of on-scalp sensors at an SNR of −20 dB to be reduced when employing a sparser array due to the projected increase in aliasing at lower sampling densities.

Results are summarised in Figure S4A. The increase in classification accuracy with increasing offsets at −5 dB when using the EBB source reconstruction approach combined with the whole brain analysis was not more pronounced than that observed for the OPM-MEG array with an inter-sensor distance of 35 mm (inter-sensor distance of 35 mm: beta = −0.423, p = 0.130; inter-sensor distance of 55 mm: beta = −0.295; p = 0.173). However, we found that accuracy increased with increasing scalp-sensor distance at −10 and −20 dB. Note that these increases were not statistically significant.

We tentatively propose that the observed advantage of moving the sensors farther away from the scalp being present also at lower SNRs when using a sparser array, may reflect a more pronounced aliasing effect at reduced sampling densities. Regarding the ROI approach, at −20 dB, we observed no decrease in classification accuracy with increasing scalp-sensor offsets at inter-sensor distances of 55 mm. Conversely, logistic regression indicated a significant decrease for sensor arrays with inter-sensor distances of 35 mm. This observation is in line with the assumption that the expected increase in aliasing at lower inter-sensor distances could provide larger scalp-sensor offsets an advantage, potentially offsetting the expected gain from placing sensors closer to cortical sources during non-invasive laminar inference.

We next repeated our simulations across scalp-sensor offsets using an OPM-MEG array with a 25 mm inter-sensor distance to examine whether the putative advantage for larger scalp-sensor offsets due to an aliasing effect diminishes with increased spatial samplings (Fig. S4B). However, we did not observe any marked changes compared to the simulations performed for an array with an inter-sensor distance of 35 mm. No significant increases or decreases were observed across scalp-sensor offsets for either classification accuracy or bias at the tested SNRs.

**Fig. S3.**
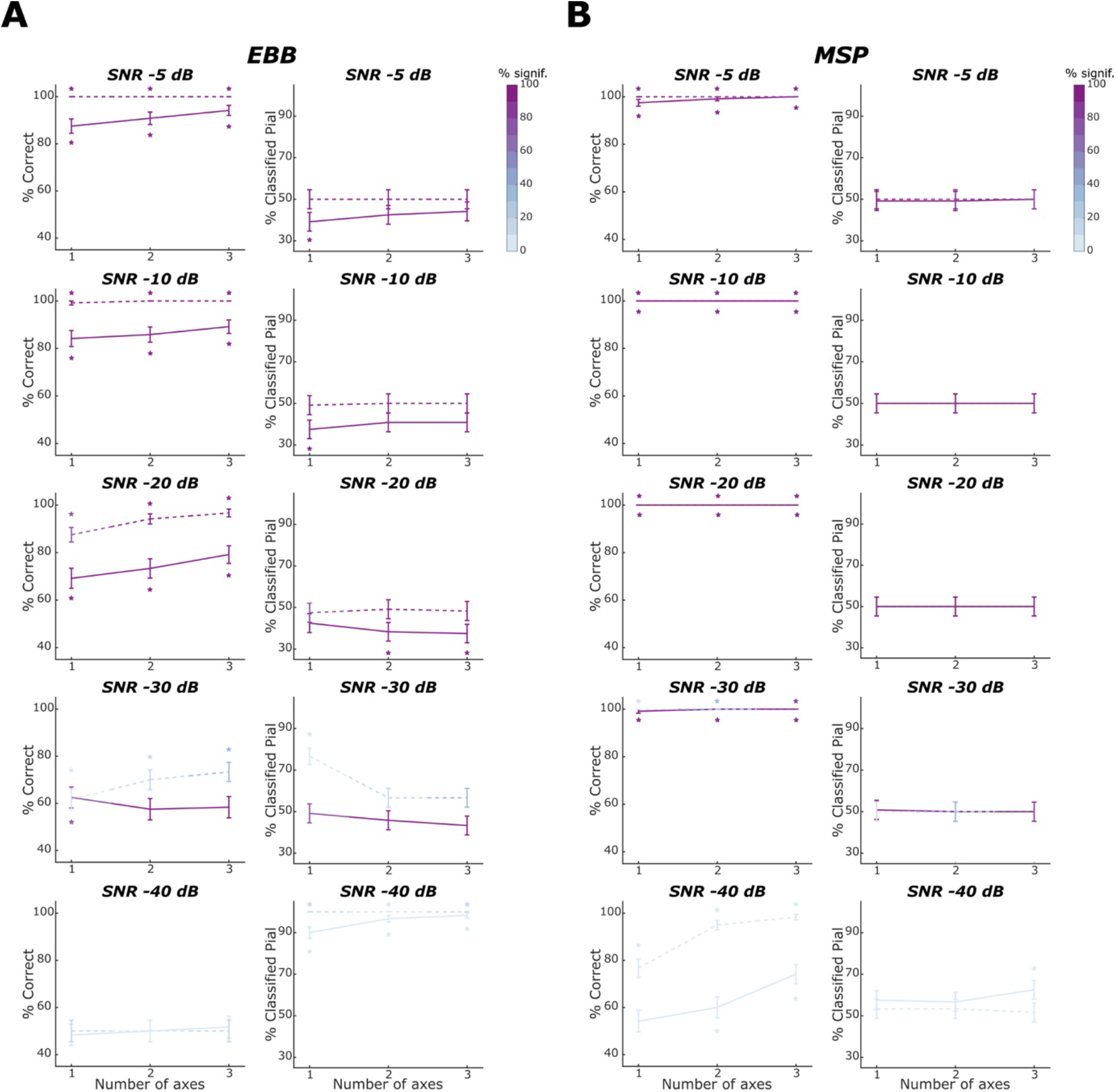
Laminar classification accuracy and bias across number of measurement axes for an OPM-MEG array with a sensor spacing of 35 mm. Solid lines denote laminar inference based on the whole-brain free energy analysis, dashed lines laminar inference based on the ROI-t-statistic analysis. Left columns within each subpanel show the percentage of correct laminar inferences, right columns show the percentage of simulations where laminar inference favoured the pial source model. SNR decreases across rows. The percentage of simulations with free energy differences or t-statistics exceeding the significance threshold is represented by the intensity of the line colour. The error bars represent the standard error. Asterisks show where the percentage is significantly above or below chance levels. **A** For the EBB approach, we found increases in classification accuracy with increasing numbers of measurement axes at SNRs of −20 dB or higher. However, these changes did not reach statistical significance. **B** For the MSP approach, classification performance was at ceiling and did not vary significantly with the number of measurement axes at all SNRs at which laminar inference was possible.

**Fig. S4.**
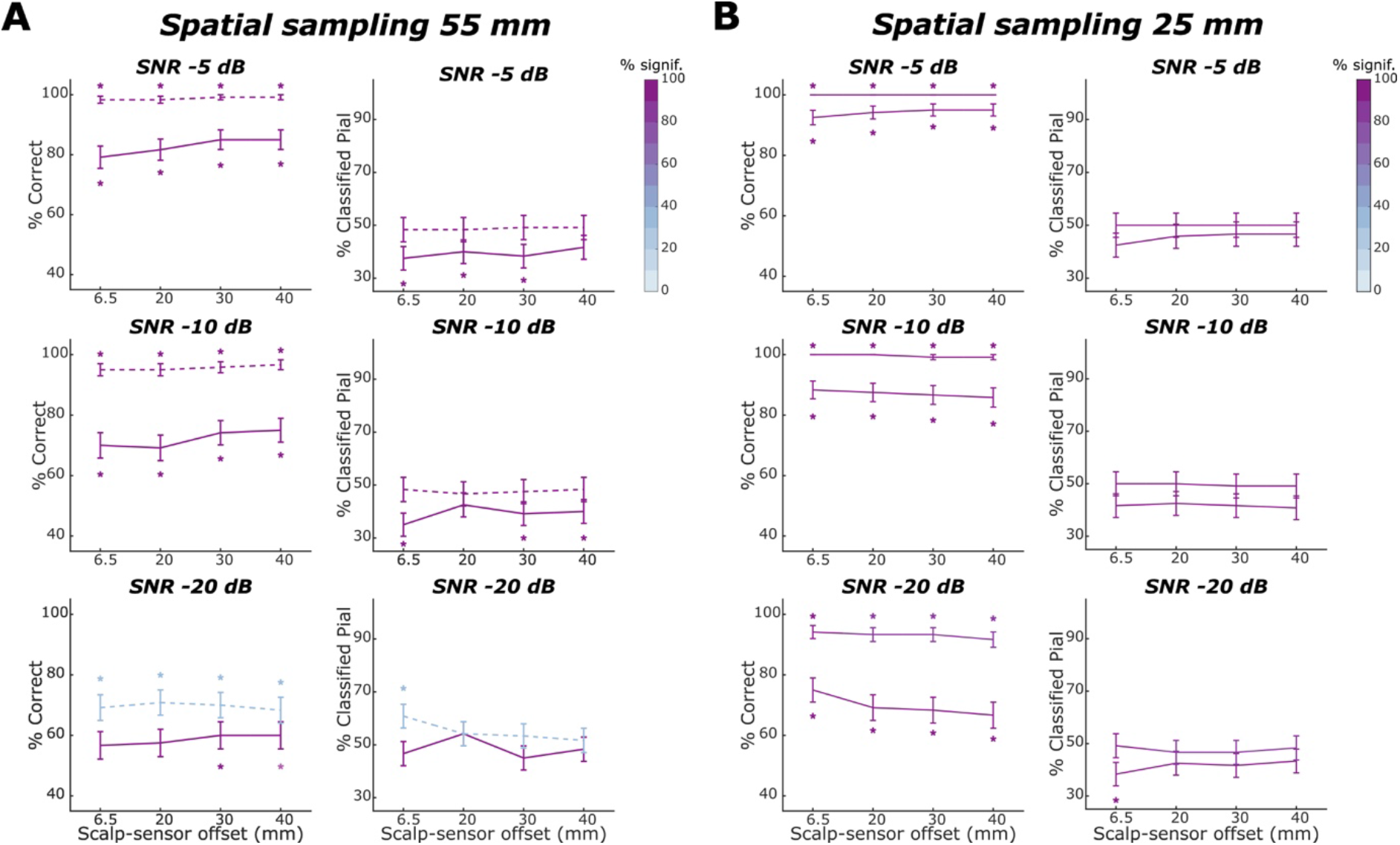
Laminar classification accuracy and bias across scalp-sensor offsets for an OPM-MEG array with a sensor spacing of 55 mm and 25 mm. Laminar inference results are shown for the EBB source reconstruction approach. Solid lines denote laminar inference based on the whole-brain free energy analysis; dashed lines laminar inference based on the ROI-t-statistic analysis. Left columns within each subpanel show the percentage of correct laminar inferences, right columns show the percentage of simulations where laminar inference favoured the pial source model. **A** For a sparser sensor spacing of 55 mm, we observed a tendency of increasing scalp-sensor distances leading to increasing laminar classification accuracy at SNRs of −5, −10 and −20 dB (all ns.). Classification bias did not vary significantly across scalp-sensor offsets. **B** For a denser sensor spacing of 25 mm, no significant changes were observed across scalp-sensor offsets for either classification accuracy or bias at the SNRs tested.

### The impact of co-registration errors at an SNR of −20 dB

We repeated our analysis on the impact of varying co-registration errors with the same simulated OPM-MEG array (inter-sensor distance = 35 mm; single axis) at a lower SNR of −20 dB (Fig. S5). For the EBB approach, we found a decrease in classification accuracy across increasing co-registration errors, which was significant for the ROI-based analysis (beta = 0.786, p <.001). For the whole-brain free energy analysis, classification accuracy was already comparatively low at zero co-registration error and the further decline in accuracy did not reach significance (beta = 0.209, ns.). Laminar inference was feasible across all co-registration errors tested, irrespectively of the analysis type employed. Classification bias remained stable across co-registration errors for the whole-brain analysis and increased with increasing co-registration errors for the ROI analysis (beta = −0.285, p = 0.050). For the MSP approach, classification accuracy was high across co-registration errors (above 90%) but decreased significantly across increasing co-registration errors for the ROI analysis (beta = 2.190, p <.01). No classification biases were observed at any co-registration error for either type of analysis. Overall, the impact of co-registration error at −20 dB resembled our observation at the higher SNR of −10 dB.

**Fig. S5.**
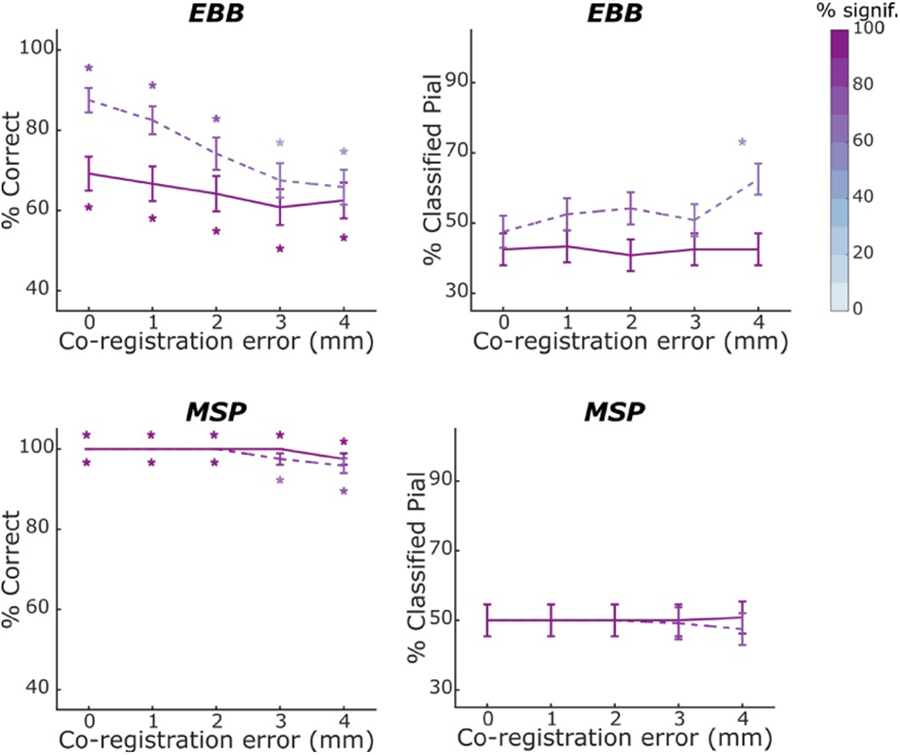
Laminar classification accuracy and bias across co-registration errors at an SNR of −20 dB. We used an OPM-MEG array with a sensor spacing of 35 mm and applied the EBB and MSP source reconstruction approaches to the simulated data. Solid lines denote laminar inference based on the whole-brain free energy analysis; dashed lines laminar inference based on the ROI-t-statistic analysis. Left columns show the percentage of correct laminar inferences, right columns show the percentage of simulations where laminar inference favoured the pial source model. For the EBB approach, we observed a decrease in classification accuracy with increasing co-registration errors. We did not observe any significant changes in classification bias with varying co-registration errors for either the whole-brain or the ROI-based analysis (B). For the MSP approach, classification accuracy decreased significantly with increasing co-registration errors. No classification biases were observed for either the whole-brain or ROI-based analyses.

